# Oops: A Language for Formalizing Fallible Biological Protocols

**DOI:** 10.1101/2025.06.17.660245

**Authors:** Matthew Shang, Eric Klavins, Gilbert Bernstein

**Affiliations:** University of Washington, Seattle, WA, USA

## Abstract

Formalizing protocols used in wetlab biological research as programs improves reproducibility by making protocols replicable and standardized. However, existing protocol languages have limited capacity for codifying error sources and standardizing error handling. When protocols inevitably go wrong, debugging must still be done by technicians, a process which is challenging for unfamiliar protocols and a scalability constraint in automated facilities. We introduce Oops, a programming language for describing biological protocols along with how they can fail. Oops makes expression of how errors affect outcomes concise through probabilistic semantics, an extensible abstraction of the lab, and metaprogramming. Given an outcome, Oops enables analyses of what errors occurred by mapping lab observations to enforced program conditions and applying existing probabilistic inference algorithms. We formalize a collection of molecular cloning protocols and present case studies demonstrating how Oops can explain errors and assess diagnostic capabilities.

## 1 Introduction

Irreproducible biomedical research has been estimated to cost $28 billion a year [10]. Although biological experiments typically report their procedures in *protocols*, it remains extremely difficult to replicate results because inadequate descriptions cause ambiguity during execution. A study in cancer biology was only able to reproduce 50 out of 193 influential experiments even after receiving clarifications from the original experiments’ authors [9]. Many of the experiments failed to replicate because they failed to match expected results over years of reproduction attempts. Lab technicians spend much of their time debugging experimental protocols because they often go wrong.

Protocols fail for a variety of reasons. The original procedure could have been confounded by an unrecognized factor, meaning that any replication attempt would be all but doomed without knowledge of that key factor. It is also possible that the technician accidentally left out a step or contaminated their workspace, or even that a random biochemical error occurred by chance. When protocols fail, it is important to determine what the cause was. A random genetic mutation is resolved by retrying the experiment, but systematic contamination requires all other experiments to be paused and lab equipment to be sanitized. To mitigate errors, practitioners have proposed protocol languages [3, 5–7, 13–15, 18, 24] that standardize biological protocols in code. Standardizing how protocols are documented reduces misinterpretation opportunities and supports robotic execution for large-scale experimentation and manufacturing.

Errors are still inevitable where biochemistry and humans are involved, despite formalized protocols and automation. However, existing protocol languages have limited if any support for specifying how protocols can go wrong. As shown by the cancer biology reproducibility study [9], figuring out what error happened is challenging for unfamiliar protocols. On the biofoundry scale, human technicians must contend with errors made in unfamiliar protocols and with outcomes confounded by uncontrolled factors, all at high volume. The troubleshooting bottleneck limits the scalability of such services all while the number of novel protocols and demand for biomanufacturing and research grows [8]. Standardizing what the errors are that could affect a protocol outcome, how they could affect it, and how to determine which error may have occurred would both assist manual and enable automated troubleshooting efforts.

The goal of existing languages is to standardize what *should* happen when executing a protocol. As a result, protocols are fixed in three ways: the outcomes of individual steps, the outcomes of the entire protocol, and the steps taken in the protocol. But when formalizing what *could* happen, all of these need to deviate. On the scale of individual steps, biophysical processes and liquid transfers have inherent continuous variance. The outcome of a protocol varies depending on both variation within steps and mistakes made that deviate from the ideal protocol. In the larger experiment, the steps taken in a protocol are purposefully modified into *controls* to check for confounding errors. Writing down protocol variants that express all these combinations of errors and controls in deterministic protocol languages is repetitive at best and intractable at worst. Moreover, even if one did describe all these variants, writing down an outcome for every possible input would be infeasible.

Our insight is that a probabilistic protocol language with transformative metaprogramming makes specifying fallible protocols with their controls extremely concise. The probabilistic framework also enables reasoning powered by traditional probabilistic inference techniques.

We design a new language called Oops that represents biological protocols with probabilistic semantics. These semantics introduce errors on two different scales: Protocols can go wrong by mistakenly running steps drawn from a discrete distribution, and steps can go wrong by drawing their quantities, such as volumes and concentrations, from continuous distributions. These erroneous steps are mapped to erroneous outcomes through models specified in a domain-specific language that abstracts the biology lab.

Probabilistic semantics alone can express fallible protocols, but the original intent becomes obfuscated. Furthermore, controls that purposefully vary a protocol still have to be written out separately. We observe that both a fallible protocol and controls are variants of the ideal protocol that can be automatically derived through transformative metaprogramming. We build a protocol metaprogramming system that separates error specification from the ideal specification and transforms protocols into control variants with user guidance.

Oops programs define distributions over experimental outcomes. Naturally, we would like to employ probabilistic inference to compute posterior distributions over errors conditioned on observed outcomes. Fortunately, observations in the lab are much like observations in existing probabilistic programming languages (PPLs). Observations in experiments can be made concurrently in control protocols running at the same time as the main protocol. Observations can also short circuit an experiment if it does not proceed as expected. Finally, observations may only happen at certain points in a protocol due to physical constraints. We develop a formalization of lab observations that casts Oops protocols as probabilistic models. We then describe probabilistic queries on these models and show how to compute them. The main contributions of this work are:

- We design a probabilistic domain-specific language that specifies both what *should* happen and what *could* happen in a biological protocol (Section 3).
- We build a protocol metaprogramming system for concisely transforming a protocol into fallible and controlled variants. (Section 4). Controls help determine what *did* happen.
- We show how to map lab observations to conditioned Oops protocols and answer probabilistic queries on protocols (Section 5).

We formalize a standard molecular cloning workflow and perform case studies of how Oops could improve practitioners’ understanding of errors in this workflow. (Section 6). Oops can not only guide an investigation into what went wrong but also help explore how a protocol can be designed to fail faster and provide more diagnostic information about failures.

## 2 Example

The polymerase chain reaction (PCR) is a workhorse protocol of modern biology [22]. To motivate and introduce Oops, we’ll show how to formalize a prototypical PCR protocol.

### 2.1 An Ideal PCR Amplification Protocol

The goal of a PCR protocol is to rapidly amplify a DNA segment of interest known as the *template*. The copying is done by DNA polymerase, an enzyme that synthesizes new DNA strands. This enzyme requires nucleotides (building blocks of DNA) and a buffer solution to operate, which are all combined into a *master mix*. Specific to the template are *forward* and *reverse primers*, which define the region to be amplified. The template, master mix, and primers are combined in a PCR machine, which repeatedly heats and cools the mixture to amplify the template.

Consider an amplification protocol in Oops:

~~~
@protocol
def amplify(F, R, T):
 M = <u>Retrieve</u>(“master_mix”)
 P = <u>Create</u>(“pcr_result”, volume=0 * uL)
 <u>Transfer</u>(M, P, 23.0 * uL)
 <u>Transfer</u>(F, P, 1.0 * uL)
 <u>Transfer</u>(R, P, 1.0 * uL)
 <u>Transfer</u>(T, P, 1.0 * uL)
 <u>React</u>(P, pcr)
 <u>Store</u>(M, P)
~~~

In Oops, which is embedded in Python, the function decorator @protocol marks a protocol. This `amplify` protocol is parameterized, which means it can be invoked from other protocols with arguments for the forward primer (`F`), reverse primer (`R`), and template (`T`).

Variables such as `F` and `M` refer to *containers* ^1^. At a given instant in time, the state of the lab can be viewed as the collective states of all containers, which we call the *inventory*. The body of a protocol is a list of *actions*, which are methods that transform an inventory. The <u>Retrieve</u> action in this protocol models a lab technician taking the `master_mix` container from a freezer and calling it `M`. Later, `M` (and `P`) is returned to the freezer using the <u>Store</u> action. Containers have an identifier and a temporary short name when on the lab bench.

After creating an empty container as `P` on the bench, liquids are transferred into `P`. Physical quantities in Oops are specified with units using the Pint [1] package. Now that `P` has all the ingredients necessary for the PCR reaction, the <u>React</u> action transforms `P` using an Oops *reaction* called `pcr`. This is a model of PCR specified by a function transforming a container that applies a basic limiting reaction model of PCR to the container’s contents:

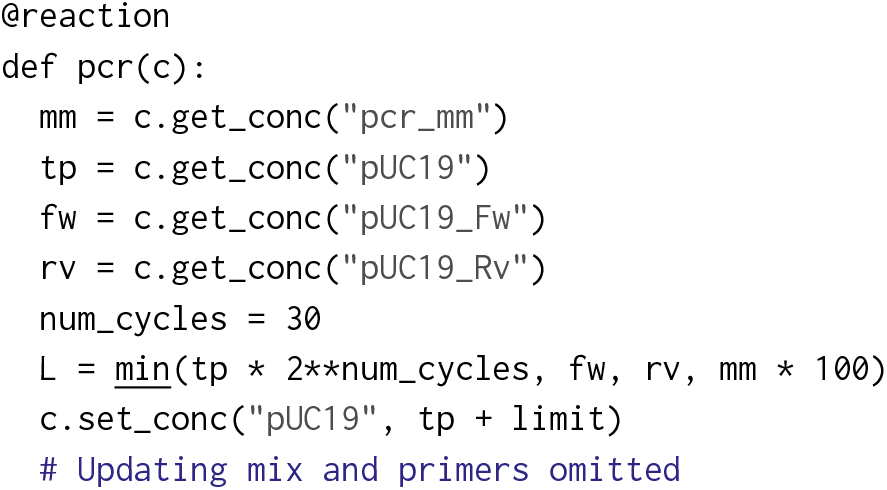

The decorator @reaction registers this function as an Oops model that can be used by protocols. The model reads the concentration (`get_conc`) of master mix (`mm`), template (`tp`), and forward (`fw`) and reverse (`rv`) primers for the pUC19 plasmid to compute a final output concentration (`L`), which is written back to the concentration of the template (`set_conc`).

While this model is simplistic, it is easy in Oops to make a model more complex when the need arises. For example, we could modify `pcr` to explicitly model each PCR heating and cooling cycle, or to work on more species than just the pUC19 plasmid by adding containers to the starting inventory.

With the generic amplification protocol and model specified, we can define a full experimental protocol that amplifies the pUC19 template:

~~~
@protocol
def experiment():
 Fw = <u>Retrieve</u>(“pUC19_Fw”)
 Rv = <u>Retrieve</u>(“pUC19_Rv”)
 T = <u>Retrieve</u>(“template”)
 amplify(Fw, Rv, T)
 P = <u>Retrieve</u>(“pcr_result”)
 <u>Observe</u>(P.pUC19 > 0.01 * mg / uL)
 <u>Store</u>(F, R, T, P)
~~~

To invoke the `amplify` protocol we defined earlier, we call it like an ordinary Python function with container bench names as arguments. The <u>Observe</u> action both specifies the expected result of this experiment and conditions the protocol on the outcome.

### 2.2 Making Mistakes

One common source of error has nothing to do with molecular biology. It is retrieving the wrong sample from the freezer. A lab technician might retrieve a forward primer for a different plasmid with a similar label in the same box in the freezer. Oops expresses errors on which step was taken with probabilistically taken branches recorded by the <u>Either</u> action:

~~~
<u>Either</u>(
 <u>Retrieve</u>(“pUC19_Fw”, F),
 <u>Retrieve</u>(“pBR322_Fw”, F),
 0.99)
~~~

The correct primer is retrieved with probability 0.99 and the incorrect primer retrieved with probability 0.01. These probabilities can be calibrated by historical data.

If every line in `experiment` was augmented with a potential mistake, the intent of the original protocol would be hard to decipher. Oops provides a metaprogramming system to separate the intended protocol from the error-prone version and apply the error rewrites automatically.

~~~
experiment = edit(experiment,
 either({“name”: “pBR322_Fw”},
    with_prob=0.01, at=“Retrieve”))
~~~

The `edit` keyword begins a transformation. This transformation only has one operation, `either`, which replaces an action with an <u>Either</u> action that sometimes runs a variation of that action. In this case, the variation is that the container name being retrieved is replaced with `pBR322_Fw`. The target location is specified with the selector syntax `at=“Retrieve “`, which selects the first <u>Retrieve</u> action. The result is an <u>Either</u> action identical to what we wrote by hand.

### 2.3 Creating Controls

A *control* is a variant of a protocol that controls for confounding factors. In our experiment, we want a *negative* control on the template to ensure that any output DNA comes from the template we put in. Negative means that we run a PCR reaction without the template. If such a negative control still amplifies the template, then we say it fails and consequently suspect contamination. Below we write out a negatively controlled version of `amplify`, with green text and red strikethroughs showing the difference from the original.

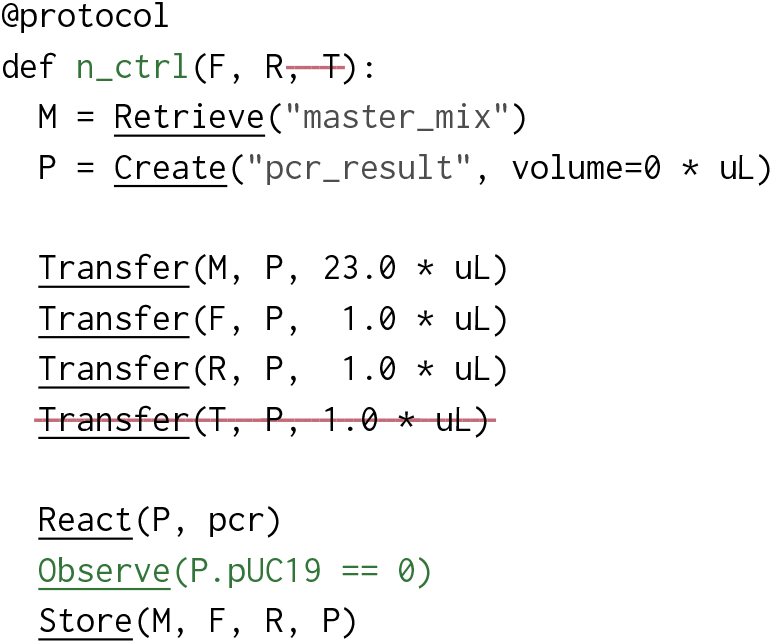

This looks similar to the original `amplify`. The template parameter is missing, along with actions that only take the template as an input. Protocol metaprogramming lets us derive the same negative control from the original protocol.

~~~
n_ctrl = edit(amplify,
 rename(“n_ctrl”),
 remove_param(“T”),
 insert(“Observe(P.pUC19 == 0)”, at=“Store(M)”))
~~~

Here, we renamed the protocol to `“n_ctrl”` and inserted an expected observation of zero output. The `remove_param` transformation removes `T` from the list of parameters and also runs a dataflow analysis to remove actions dependent on `T`. Also note the selector syntax `“Store(M)”`, which selects a location inside the protocol to perform the edit. In this case, `“Store(M)”` looks for the first <u>Store</u> action with an `M`. See Section 4 for more details, including on how we also derive positive control variants.

Finally, we use more metaprogramming to insert this control into the experimental protocol right before the invocation of `amplify`:

~~~
experiment = edit(experiment,
 run(n_ctrl, at=“amplify”, args=(“Fw”, “Rv”)))
~~~

Again, we used a selector `“amplify”` that targets the invocation of the `amplify` subprotocol.

This amplification procedure is just the first part of the molecular cloning workflow we formalize in the evaluation (Section 6). There, we also demonstrate how Oops can determine how helpful this negative control is for figuring out the wrong primer error.

## 3 The Oops Language

In this section, we define Oops formally, with the goal of using it to specify what *should* happen in a biological protocol and, just as importantly, what *could* happen. We’ll cast the lab as a computing platform by distilling representations of data—containers (3.1) and inventories (3.2)—and of code that manipulates this data—actions (3.3), protocols (3.4), and reactions (3.5).

### 3.1 Containers

*Containers* generalize test tubes, cuvettes, flasks, Petri dishes, and other vessels designed to hold, manipulate, and analyze biological materials. On the lowest level, Oops models containers as dictionaries keyed by *properties*, which are strings. Examples of properties include volume, DNA concentration, and number of cell colonies.

Oops makes a distinction between *mixtures*, which represent liquid mixtures with a volume and concentrations of species, and non-mixtures. Liquid transfer is not allowed from non-mixtures to mixtures. In the implementation, containers expose a basic API allowing get and set operations while mixtures extend this interface to track concentration-valued properties and volume.

### 3.2 Inventories

An *inventory* represents a snapshot of the lab and consists of containers, observations that have been made, and other ambient properties, such as room temperature. Containers in an inventory can be stored in a freezer, or out on the workbench. Each container is identified by a full name and a short bench name that is temporarily assigned when placed on the bench during a protocol. For example, a test tube with a plasmid might be stored in a freezer with full name “pUC19-GFP-20240929”, retrieved and referenced as simply “pG” in a bacterial transformation protocol, and then stored back in the freezer. Both full names and bench names exist in a global scope. Containers can be retrieved multiple times and even when on the bench, but may only be stored when on the bench.

### 3.3 Actions

An *action* is a method that represents a lab operation and describes how to transform an inventory with that operation. Actions are instantiated by parameters that dictate how the action behaves when called on an inventory, such as which bench name it should use or how much volume should be transferred. Actions are focused on producing results that can be interacted with, so Oops is not concerned with intermediate stages that are unobservable or that do not matter for the modeled set of errors and outcomes.

Figure 2 lists a core set of actions included in Oops. To give a flavor of how actions describe lab operations and how they can be composed into protocols, we describe them in more detail next. All actions must implement an interface that takes an inventory and returns an inventory.

**Figure 1.**
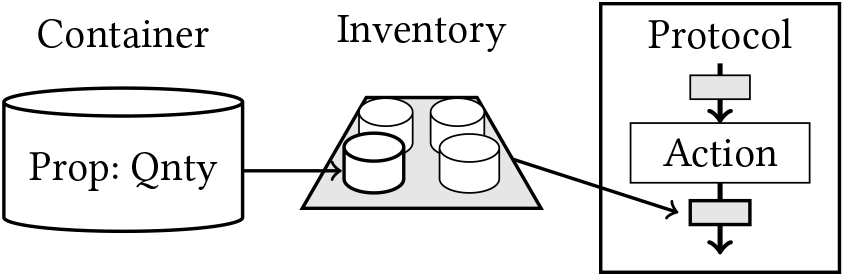
Protocols, actions, inventories, and containers.

**Figure 2.**
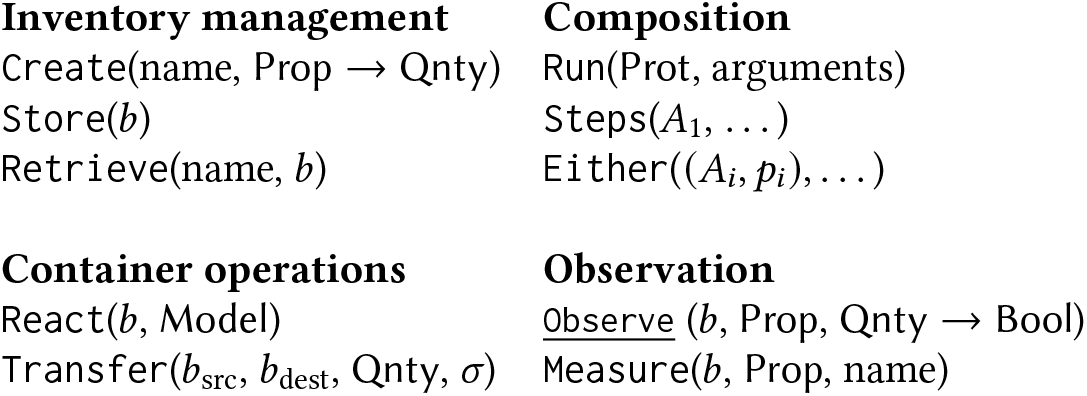
Actions in Oops. *b*’s are bench names, *A*’s are actions, *p*’s are probabilities, and *σ*’s are standard deviations.

#### Inventory management

A surprisingly common error made in a biology lab is mislabeling or misreading containers, which is less surprising considering that storage tubes can be tiny, labels be hand written, and hundreds of tubes packed into a single box, hundreds of boxes packed into a single freezer, and dozens of freezers supporting a large lab. The <u>Create</u>, <u>Store</u>, and <u>Retrieve</u> actions express the management and mismanagement of containers.

Specifically, <u>Create</u> models the use of a fresh container that has been given a full identifier and placed on the bench. The <u>Retrieve</u> action represents retrieving a container from the lab store and placing it on the bench, while <u>Store</u> represents storing a container currently on the bench back into the store. Containers can be retrieved again once already on the bench, but can only be stored when on the bench and cannot be stored again without being retrieved first.

#### Composition

The <u>Run</u> action runs an entire *protocol*, which we discuss in Section 3.4. A key action is <u>Either</u>, which is a nondeterministic branch. It is instantiated with a list of actions *A*_*i*_ and a list of probabilities *p*_*i*_ that sum to one. When called on an inventory, the <u>Either</u> action chooses one of its *A*_*i*_ to execute with probability *p*_*i*_. The <u>Steps</u> action allows for a sequence of actions to happen as a unit in <u>Either</u>.

The main purpose of <u>Either</u> is to express errors that can be made when executing a step, such as a mistaken retrieval. The probabilities *p*_*i*_ encode a prior on these mistakes.

#### Container operations

Both <u>React</u> and <u>Transfer</u> model transformations of containers. The <u>React</u> action runs a model of a reaction on a container. We defer discussion of how Oops expresses models to Section 3.5.

The <u>Transfer</u> action transfers a volume of liquid from one container to another. When pipetting in the lab, the actual volume transferred depends on the user’s skill and can be modeled as locally Gaussian centered at the desired volume with standard deviation *σ* corresponding to skill. When transferring small volumes, fluctuations can matter. What *should* happen is recovered by setting *σ* = 0.

To model the transfer of *t* units of liquid from container *A* to container *B*, we first set *u* = *t +* normal (0, *σ)* and clamp *u* to be between 0 and the volume of *A*. Let *𝓋*_old_ (*X)* be the volume of *X* before the transfer and *𝓋*_new_ (*X)* be the volume after. The new volume of *B* is simply *𝓋*_new_ *B* = *𝓋*_old_ (*B) +u*, while the new volume of *A* is *𝓋*_new_ *A* = *𝓋*_old_ (*A) − u*. For any concentration property *p* in either *A* or *B*, let *c*_old_ (*X)* be the concentration of *p* in *X* before the transfer and *c*_new_ (*X)* be the concentration after. Then the new concentrations of *p* are

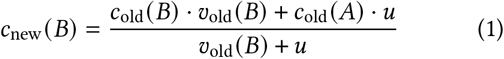

and *c*_new_(*A*) = *c*_old_ (*A*).

Since Oops distinguishes between mixture and non-mixture containers, transfers must have a mixture as a source container. However, non-mixture containers can be destinations to model operations like transferring contents into a machine for analysis. For non-mixture destinations, we assume the existing volume to be zero.

#### Observation

An inventory includes measurements in its state, modeling a lab technician’s lab notebook recording a trace of their experiment. The <u>Measure</u> action records the quantity associated with a property with a label so that it can be retrieved when analyzing a protocol trajectory.

<u>Observe</u> formalizes an expectation of what a property should be. It abstracts measurements that serve as verification steps that may short-circuit and halt the entire experiment. It also represents measurements taken in controls. In addition, <u>Observe</u> allows for conditioning, which we discuss further in Section 5. Its interface takes a container, a property *p* of that container, and an expected predicate *f* for that property. When the Oops framework executes an <u>Observe</u> (that is not in a control protocol), the program halts if the value of *p* does not satisfy *f*.

Both <u>Measure</u> and <u>Observe</u> abstract various measurements that can be taken in a lab, whether by eye, a machine, or an instrument. As an analogy, <u>Observe</u> actions are assertions, while <u>Measure</u> actions are print statements.

### 3.4 Protocols

A *protocol* is a list of actions that are executed sequentially. Oops enables composition and reuse of protocols through parameterized protocols, which means that protocols can be defined with container-valued parameters that are substituted for arguments at runtime. The <u>Run</u> action keeps track of arguments. Protocols have names and can be marked as controls, which distinguishes control protocols running concurrently from the main protocol.

We implement the Oops language as an embedded DSL in Python. The @protocol decorator parses a Python function AST into an Oops protocol AST, which is really a tree of actions. To connect action definitions to action instances created in the DSL, Oops provides an @action decorator that registers an action as a keyword and generates parsing code.

### 3.5 Reactions

To specify what outcome results from each path of actions taken through a protocol, Oops needs a model mapping inputs to an outcome. Otherwise, the user would have to specify the outcomes of combinatorially many possible paths, each representing a different subset of mistakes. So far, the protocol language as described implements part of this model. Containers and actions like <u>Transfer</u> model the evolution of mixtures of species by tracking total volume and concentrations inside the mixture. However, nothing in this framework encodes how those species interact, which is the reason why anything interesting at all happens in the lab.

Oops allows the user to define the most interesting phenomena of the model, such as chemical reactions and cell growth, with *reactions*, which are functions that transform a single container to its state after a biological or chemical process runs to completion. Reactions are registered using the @reaction decorator and invoked using the <u>React</u> action. Since many processes run to completion before the next interaction with the container, usually, modeling the results of a reaction is sufficient. We purposefully do not aim to model interactions in full biophysical detail, since many biological processes do not have generic mathematical theories and modeling on first principles leads to infinite regress.

Exposing parts of the model definition to the user enables them to encode their own inductive reasoning about complex biological processes. Once the Oops model no longer reflects lab observations, the reactions can be updated to encompass new findings. This incorporates the *phenomenological* theories found in empirical lab work that combine with theories of underlying mechanisms to explain how outcomes come from inputs.

## 4 Protocol Metaprogramming

The goal of this section is to derive protocols that specify what could happen and determine what did happen from a protocol that describes what should happen. To that end, we show how program transformation enables transforming protocols into positive and negative controls (Section 4.1) and separating the specification of what ideally happens from erroneous deviations through transformations (Section 4.2). We then build a metaprogramming language to express these transformations concisely (Section 4.3).

### 4.1 Metaprogramming Controls

There are two main types of controls. *Positive* controls confirm that experimental conditions are capable of producing results. *Negative* controls support hypotheses that results are due to a given experimental variable. Together, they separate the experimental outcome from confounding factors and verify that the setup works as intended. Both provide valuable information for determining what happened when a protocol goes wrong.

Given a protocol *p* and an experimental variable *X* in the protocol, running a negative control on *X* means executing *p* with a null version of *X*. Examples include omitting *X* in a reaction or using an inert substance in place of *X*. Running a positive control on *X* means executing *p* with other experimental variables fixed to values that are known to produce a positive result. In a PCR protocol, a negative control on the template means running a PCR without any template DNA to check for contamination, while a positive control on the primers fixes the template to a sequence the primers are known to amplify to check that the primers work.

We discovered that these methodological patterns correspond to systematic transformations of code. Given an Oops protocol with some parameters, a negative control can be derived from the original by removing a parameter representing an experimental variable, eliminating actions dependent on that parameter along with actions dependent on those, and appending an observation of a negative result. Similarly, a positive control can be derived by replacing parameters outside of parameters to be controlled with constant values and appending an observation of a positive result. Figure 3 shows an example of these transformations on a simple protocol.

**Figure 3.**
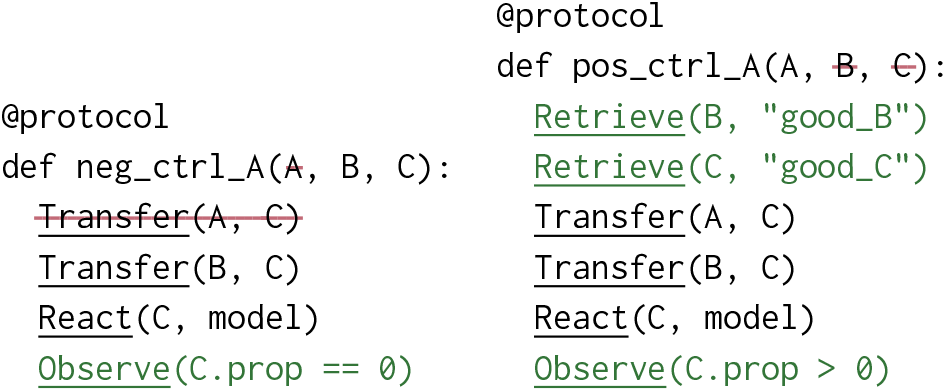
Transforming a protocol into (left) a negative control and (right) a positive control. (Left) removing parameter `A` also removes the <u>Transfer</u> action using `A` as a source. (Right) `B` and `C` are specialized to known containers.

### 4.2 Encoding Errors with Metaprogramming

Protocol transformations also enable a workflow that allows users to separate the ideal protocol from that protocol with its controls and errors. This preserves the intent of the original protocol and makes it easy to iterate on both the protocol and its errors. Since controls are derived from the original protocol, changes to the original are automatically reflected in the controls. This is in the same spirit as user-schedulable languages [11, 17] except that transformations that add errors do not preserve semantics.

### 4.3 The Oops Metaprogramming Language

To express and compose code transformations, Oops exposes metaprogramming primitives that operate on protocols. Table 1 describes some of the primitives available. Sequences of transforms are applied to protocols by using the `edit` function, which has the interface `edit(protocol, ops…)`. This makes it easy to group related transforms and inspect intermediate programs.

**Table 1.** Some of the metaprogramming operations in Oops.

| Transform | Description |
| --- | --- |
| `remove_param(x)` | Remove $x$ from the parameters and perform a forward dataflow analysis [4] to remove dependent actions. |
| `specialize(x, v)` | Remove $x$ from the parameters of $p$ and replace all $x$'s with $v$. |
| `insert(a, at)` | Insert action $a$ before the first action matching $at$. |
| `replace(a, at)` | Replace the first action matching $at$ with $a$. |
| `either(a, at, prob)` | Replace the first action $a'$ matching $at$ with `Either(a', a, prob)`. |
| `run(p, at, args)` | Insert protocol $p$ invoked with $args$ before the first action matching $at$. |
| `rename(n)` | Rename to $n$. |

To specify which action a transformation should target, we build a selector syntax. A selector with just the name of an action, such as `“Store”`, finds the first top-level <u>Store</u> action in a protocol. If there are multiple matching actions, a selector can be further specified with arguments in the action (such as a container name) and the names of any number of containing protocols. For example, a <u>Store</u>(A) action located in a subprotocol named `assemble` can be referenced with `“assemble(Store(A))”`.

## 5 What Happened?

Recall that <u>Observe</u> actions function as assertions in protocols. We view these assertions as generalizations of two types of observations made in the lab while running protocols (Section 5.1). The joint distribution over errors and assertions leads to an interpretation of Oops protocols as probabilistic models (Section 5.2). We then define queries on these models motivated by questions about reproducibility (Section 5.3) and show how to compute these queries (Section 5.4).

### 5.1 Unifying Observations

In a software debugger, the entire state of the program is visible after every operation. In the lab, it is not possible or practical to exactly measure every physical property at every point. We focus on two main kinds of observations one might make while running a protocol:

#### Sanity checks

Between some steps in a protocol, it is sensible to measure a quantity that confirms if the protocol is going as expected. For example, after amplifying template DNA using a qPCR machine, gel electrophoresis measures the concentrations of DNA fragments at different lengths. If the concentration at the length of the target DNA is nonexistent or low relative to other lengths, then something has gone wrong and it is not worth continuing forward.

#### Controls

Other than the main protocol, a protocol might have subprotocols that are run concurrently (in the case of controls) or later (when troubleshooting). These protocols also result in observations that give meaningful information about the original protocol.

Both kinds of observations have pass/fail outcomes. What counts as a success or failure is determined by a property and a predicate that defines acceptable measurements of that property. In addition, both kinds of observations serve as breakpoints. A failed sanity check halts execution of the protocol, while a failed control observation provides evidence for a confounding factor. As a corollary, the knowledge that a sanity check occurred provides extra information that all previous sanity checks must have succeeded. Thus, the <u>Observe</u> action can represent both kinds of observations.

### 5.2 Oops Protocols as Probabilistic Models

An Oops protocol is a probabilistic model; a distribution over possible experiment outcomes. The random variables are discrete errors *E*_*i*_, which come from <u>Either</u> actions, control assertions *Y*_*i*_, which come from <u>Observe</u> actions in controls, and sanity check assertions *Z*_*i*_, which come from <u>Observe</u> actions in the protocol. Figure 4 visualizes an example. An Oops protocol defines a joint distribution over all possible experiment outcomes

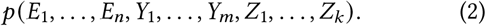

**Figure 4.**
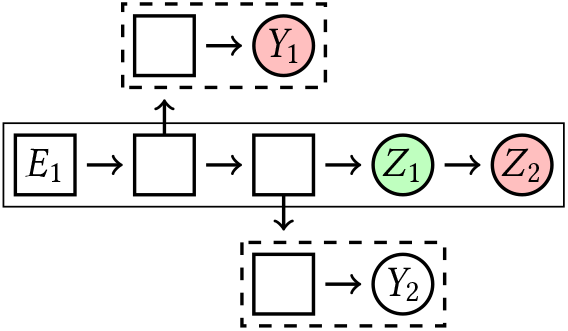
An Oops protocol viewed as a probabilistic model. The large solid box is the protocol and dashed boxes indicate controls. There is one potential error at *E*_1_. Knowing that the main protocol failed at *Z*_2_ determines that observation *Z*_1_ succeeded. The control observation at *Y*_1_ is known to have failed, but the control result *Y*_2_ is unknown.

The domain of error *E*_*i*_ depends on the possible mistake actions and includes the possibility of not making a mistake. The domains of the *Y*_*i*_ are {T, F, ⊥}, where ⊥ represents an unobserved value. Since *Z*_*i*_ failing implies that every previous sanity check *Z* _*j*_ with *j < i* must have passed, we represent the outcome of the main protocol with a single random variable *Z* whose domain is over the sanity checks along with a value T that indicates the entire protocol succeeded. In this way, every trace of an Oops protocol has values defined for every variable.

### 5.3 Motivating Probabilistic Queries

Given the viewpoint of protocols as probabilistic models, we now pose probabilistic queries on those models that are motivated by questions about a protocol’s reproducibility.

#### Error explanation

Given a protocol, we would like to explain what went wrong. Knowing that a protocol went wrong means that some sanity check failed, so our query is for a conditional distribution over each error given the first failed check. We may also know the results of some controls depending on which were run either during or after the experiment. This leads to the posterior distribution

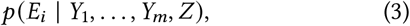

where known control results *Y*_*i*_ are set to T or F and unknown results are set to ⊥.

#### Mutual information

Given an error, we would like to know if a control helps to explain the error. We can quantify the helpfulness with the *mutual information* between *E*_*i*_, the error, and *Y*_*j*_, the control outcome. From here, we omit the subscripts for clarity. The mutual information *I (E*; *Y)* is defined as the difference in *entropy* between the prior of *E* and the posterior of *E* conditioned on *Y*; that is, *I* (*E*; *Y*) = *H* (*E*) − *H* (*E* | *Y*). In turn, the entropy of a discrete distribution *X* is defined by *H* (*X*) = Σ_*x*∈𝒳_ *p*_*X*_ (*x*) log *p*_*X*_ (*x*). Entropy quantifies the amount of information needed to describe the state of *X*. If conditioning an error on a control result leads to a distribution with less entropy than the original posterior, then the control provides information about that error.

To quantify the helpfulness of a control over all possible outcomes of a protocol, we consider the *conditioned* mutual information *I (E*; *Y* | *Z)* where *Z* is the distribution over outcomes [27]. Recall that the domain of outcomes is all sanity check observations, where a failed observation halts the entire experiment. It can be shown that for jointly discrete random variables *E, Y*, and *Z*, Equation 4 holds.

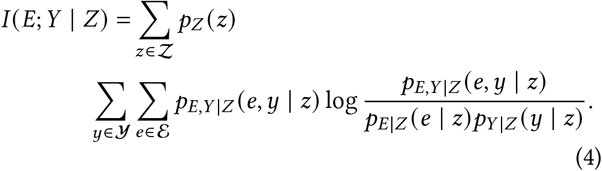

### 5.4 Computing Probabilistic Queries

All of the distributions of interest are manipulations of the joint distribution. In particular, we need to support marginalization and conditioning [23]. To compute these distributions, we use a sampling-based approach. This algorithm does not scale efficiently in the number of errors introduced to the protocol but is sufficient for our case studies.

In a preprocessing step, we collect an empirical distribution 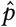 by sampling the protocol *p*. Mistakes can be rare and multiple rare mistakes might have to occur for a failure outcome. As a result, we enumerate all choices of error possibilities and sample a separate empirical distribution for each. Let 𝒞= ℰ_1_ × … × ℰ_*n*_, where ℰ_*i*_ are the possibilities made in error *E*_*i*_. For every choice of errors and non-errors *c* ∈ *𝒞*, we fix a version of the protocol *p*_*c*_ that always and only makes the mistakes in *c*. Then, we collect an empirical distribution 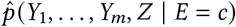 by collecting *S* samples 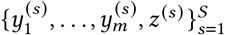. For clarity, we use *E* = *c* as shorthand for the full assignment of each *E*_*i*_ to the choice *c*_*i*_.

To answer queries, we provide a generic recipe for marginalization and conditioning.

#### Marginalization

Suppose that we want to marginalize out all variables other than *Z*. First, we sum over all enumerated choices:

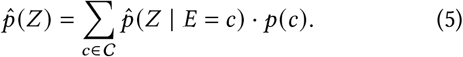

The weight *p (c)* can be calculated from the prior on the *E*_*i*_ defined by <u>Either</u> actions in the protocol. The marginalized distribution for a given choice is

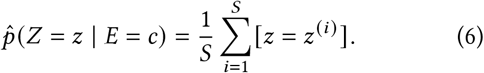

The process is similar for other marginalizations; we just keep the samples that agree in the remaining variables.

#### Conditioning

Suppose that we want to condition on *Z* and we’re interested in the posterior of one error *E*_*i*_. By definition of the conditional distribution, we have

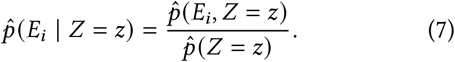

Now, we just marginalize the joint distribution.

## 6 Evaluation

We formalize an end-to-end molecular cloning workflow. Using Oops, we answer three questions about the cloning protocol motivated by our original problems:

A. What went wrong?
B. Does a control help find an error?
C. How quickly can an error condition be discovered?

### 6.1 Formalized Molecular Cloning

The goal of molecular cloning is to assemble DNA *molecules* and *clone* them into a population of cells [21]. First, in the construction phase, the DNA to be cloned is amplified by PCR and combined with vector DNA to form recombinant DNA molecules. Next, in the transformation phase, specially treated bacteria are induced to take up these molecules and replicate them. These bacteria replicate exponentially to generate a large number of copies of the original molecule. Finally, in the verification phase, the molecule is extracted from the cells and verified to have the DNA of interest.

Figure 5 shows a high level view of the workflow we use to evaluate Oops in this paper. The construction phase consists of two protocols. An amplification protocol like the one in Section 2 amplifies DNA coding for the green fluorescent protein (GFP), which is the DNA to be cloned, and a plasmid vector known as pUC19. A *plasmid* is a small, circular DNA molecule that replicates independently of a cell’s chromosomal DNA. Then, a Gibson assembly protocol assembles the plasmid with GFP to produce recombinant molecules.

**Figure 5.**
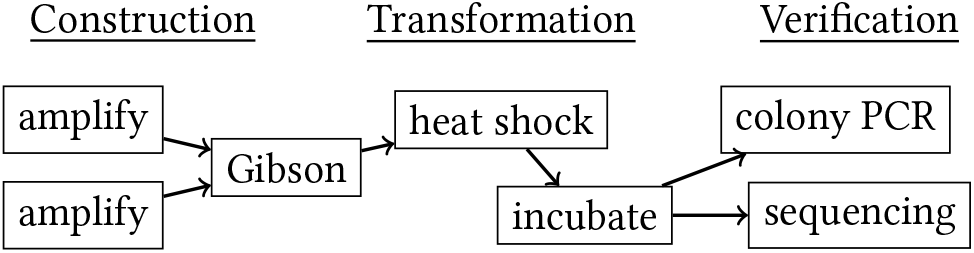
A flowchart of the molecular cloning protocols.

The transformation phase consists of a heat shock protocol and an incubation protocol. During heat shock, specially treated *E. coli* are suddenly heated and cooled to take up the recombinant plasmids. The cells are plated onto agar plates and incubated overnight to multiply. Since not every cell takes up a plasmid, transformed cells are selected for by adding an antibiotic to the plates that the pUC19 plasmid codes resistance for.

Finally, a technician selects a colony on the plate and scrapes it off for verification that the cells replicated the correct recombinant molecule. This involves two protocols: a colony PCR protocol, which is a quick verification that the lengths of replicated DNA match what is expected, and a sequencing protocol, which is expensive but checks that the DNA exactly matches what is expected.

We also introduce the errors shown in Table 2 and control protocols shown in Table 3.

**Table 2.** Errors introduced to our protocol.

| Error | Description |
| --- | --- |
| Primer | Wrong or badly designed primer |
| Reagents | Degraded PCR master mix and pUC19 forward primer are used in an amplification |
| Assembly | Misassembly during Gibson assembly, including flipped inserts |
| Antibiotic | Wrong or missing antibiotic plated |
| Mixing | Harsh mixing breaks cells |
| Contam. | Competent cells contaminated with plasmid |

**Table 3.** Controls available in our protocol.

| Control | Description |
| --- | --- |
| +GFP | Positive control on GFP amplification |
| +Vector | Positive control on pUC19 amplification |
| -Plasmid | Transform cells without adding plasmid |
| +Cells | Positive control on cell transformation with known plasmid |

### 6.2 Explaining What Happened

An early sign something has gone wrong in a molecular cloning pipeline is that the number of colonies on the plate incubated overnight is lower than expected. We use Oops to build an interactive “debugger” for the molecular cloning model that could guide a technician towards where to look. The debugger can be queried with a failed observation:

~~~
>>> debugger.explain(“xform(Observe(Plate))”)
Explanations for 25 <= colonies <= 150 in Plate:
 Error 1:
   Degrade(“Fw”, “pUC19_Fw”, …): 0.75
   Skip(): 0.25
   … (remaining errors omitted)
~~~

Explaining a failed observation means conditioning the protocol at that observation and inspecting the posterior distributions on errors. Here, we’ve conditioned on an <u>Observe</u> measuring the number of colonies in a container labeled `Plate`. In return, we get the marginal likelihood of the choices in every error, reported in decreasing order. According to the model, we might want to first check for degraded reagents that would cause low efficiency amplification and subsequently low efficiency transformation. The “control unknown” chart in Figure 6 visualizes initial conditional likelihoods for three errors.

**Figure 6.**
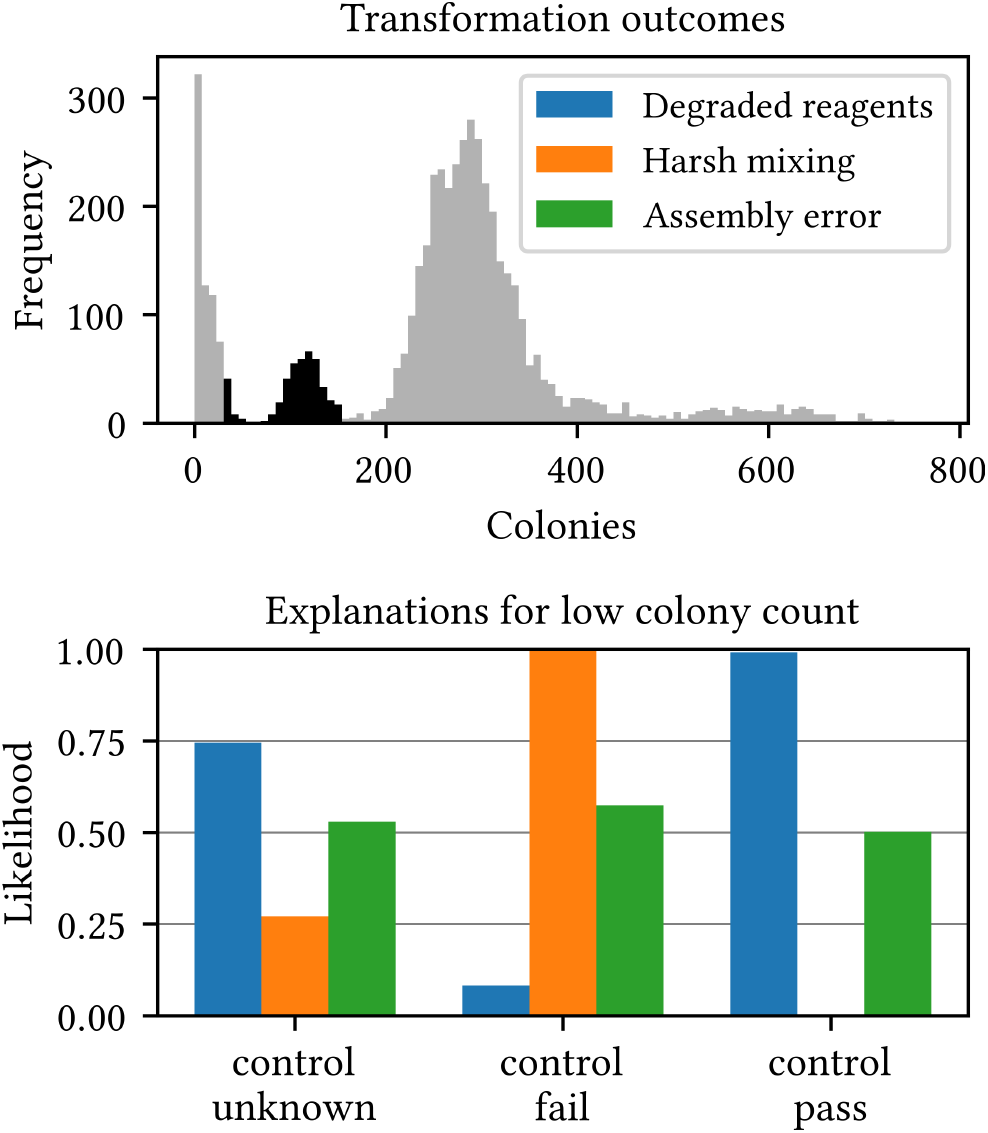
Top: Colony count outcomes of the bacterial transformation stage. Bottom: explaining three types of error conditioned on observing a low number of colonies (highlighted region in the top) and the outcome of a control.

Now suppose that during the protocol, a positive control on the competent cells during transformation was run (this is called “+Cells” in Table 3). This control attempts to transform cells with a plasmid known to work well, checking that the cells can be transformed in the first place. If the positive control passes but the transformation did not work well, then there must be an issue with the actual plasmid being cloned. On the other hand, if the positive control fails, then something must have happened to the cells.

Suppose the positive control passed. We can ask the Oops debugger to condition on the control succeeding:

~~~
>>> debugger.condition(
       “xform_pos_p(Observe(Plate))”, True)
>>> debugger.explain(“xform(Observe(Plate))”)
Explanations for 25 <= colonies <= 150 in Plate:
   Error 1
     Degrade(“Fw”, “pUC19_Fw”, …): 1.00
    …
~~~

The model suggests that the reagents must have been degraded, leading to low transformation efficiency even though the control passed. Similarly, we can condition on a failed positive control. Figure 6 visualizes how conditioning on the positive control result affects the likelihoods of different errors. By adding more known observations and ruling out errors, the Oops debugger can guide a technician in their investigation of what went wrong. If all errors in the model are accounted for, then the actual cause of failure must be outside the model.

### 6.3 Information Gain from Controls

A control provides a baseline to compare the primary experimental results against. Ideally, a control should eliminate alternate explanations so that it is certain that observed effects are due to a manipulated variable. Since controls are usually executed concurrently with the main protocol, one design question asks if it is worth running a control. Given an error of interest, we can quantify how helpful a control is for figuring out if the error happened by studying the *mutual information* between the random variable that the error happened and the random variable that the control passed or failed. We define mutual information formally in Section 5.3. Informally, mutual information quantifies the information gained about the error given the control result.

For example, focusing on posteriors for degraded reagents in Figure 6, conditioning on either control result concentrates more mass towards either the reagents being degraded or not being degraded, corresponding to information gain.

We use Equation 4 to compute conditional mutual information between every error and control implemented in our molecular cloning protocol (in tables 2 and 3). Figure 7 visualizes the results. It suggests that every control contributes bits of information towards every error other than misassembly.

**Figure 7.**
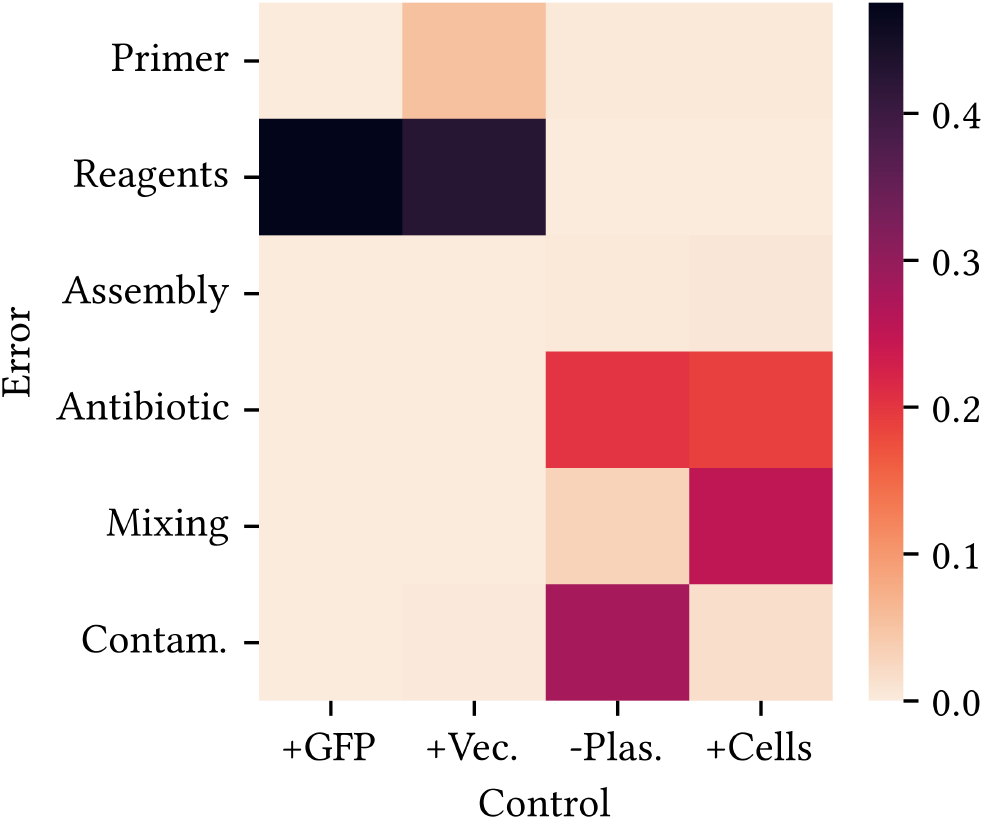
Conditional mutual information visualized for every pair of error and control. A dark color for a given error and control indicates high expected information gain in the error when the control outcome is known. A bright color means that the error and control are largely independent.

### 6.4 Failing Fast

In large scale experimental workflows executed in industrial biofoundries, massively parallel cloning processes may take more than a week to complete, at significant cost. It is beneficial when executing this protocol to know that something has gone wrong before completing the entire process, and the earlier the better. If the protocol is going to fail, it better fail fast. One way to fail faster is to add verification steps, such as controls, at the cost of making the entire protocol longer on successful runs. We analyze the impact of adding two different verification steps to our molecular cloning model to see how it could fail faster.

One step is to verify that the lengths of amplified DNA match the lengths of what is expected when amplifying the vector and insert. Each piece of DNA has a length measured in the number of base pairs making up the strand. In a protocol called gel electrophoresis, the output from a PCR is placed in a gel and molecules of different lengths are separated by an electric current. This checks that the amplification reaction worked as expected.

Another verification step we explore is a set of controls on the bacterial transformation. These are the “-Plasmid” and “+Cells” controls described in Table 3. This set of controls checks that the bacterial transformation works as expected.

Using Oops metaprogramming, we create two variants of the original molecular cloning protocol with the verification steps inserted. Then, we simulate 10,000 runs of each conditioned on a failed end result to collect distributions over the first failed observation (shown in Figure 8). The progress at an observation *0* is defined by

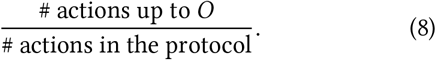

**Figure 8.**
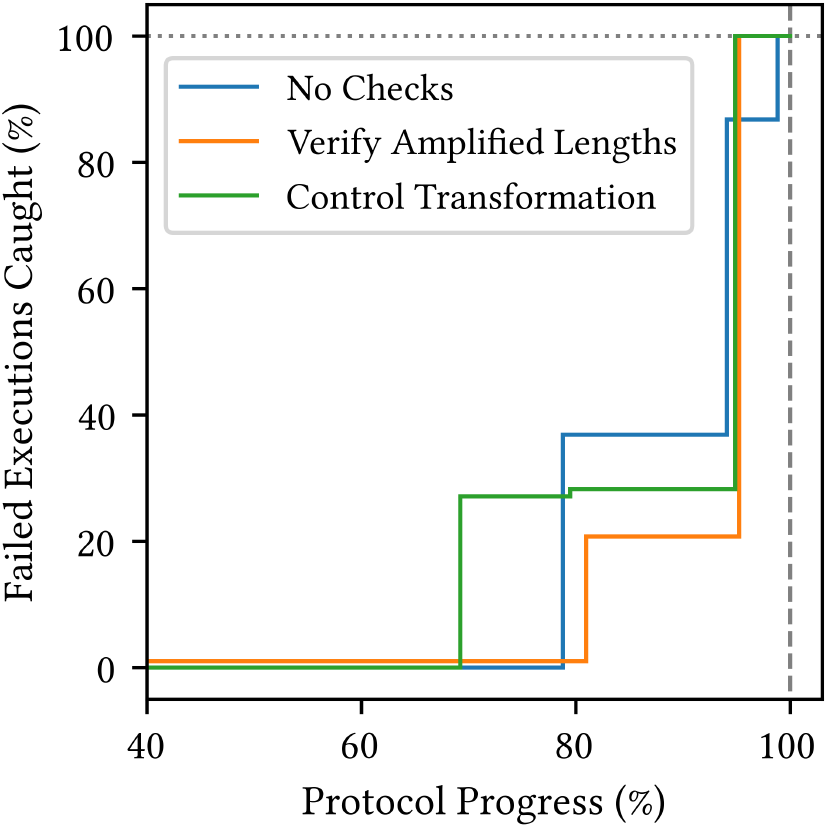
Where and how likely three molecular cloning protocol designs are to fail first, given that they ultimately fail to clone the insert.

Combinator actions such as <u>Either</u> and actions run in control protocols are not included in these counts.

Introducing the gel step fails a small number of executions early in the protocol, as shown by the orange line entering the plot (Figure 8) at a nonzero height. However, the gel introduces extra work before the rest of the protocol can be finished. So, the next step that can fail occurs later in the protocol. According to the Oops model, a more effective method of failing fast are the controls on transformation, represented by the green line. Since transformation controls can be run concurrently with the original transformation, this protocol fails faster.

## 7 Related Work

### Protocol Languages

Previous protocol languages focus mainly on standardizing what should happen in the protocol. On one end, languages compile to natural language descriptions meant for humans to execute, such as in protocols.io [20] and BioCoder [5]. On the other end, protocols are formalized to a level of detail that allows robots and microfluidic devices to execute them, such as Autoprotocol [13], Opentrons [14], and Puddle [26]. Other languages such as BioScript [15] and LabOP [6] have representations for both humans and machinery to execute. The key contribution of Oops is to introduce another axis of formalizing what else could happen when running a protocol.

Abate et al. [3] and followup work by Cardelli et al. [7] design a language with probabilistic semantics to characterize uncertainties. While they propagate stochasticity through a deterministic sequence of protocol operations, our language makes the operations executed stochastic as well. However, their Gaussian semantics enable an efficient estimator for gradients through simulated protocols.

### Automation Systems

Lab automation systems such as Aquarium [24], Benchling [2], and Synthace’s Antha [18] combine interfaces for standardizing protocols with electronic notebooks for recording data. They also schedule operations to be executed by humans or robots, forming an “operating system” for the lab. To the best of our knowledge, existing systems focus on standardizing what should happen in protocols. While historical execution data recorded in these systems can be mined for historical failure reasons, their standardizations do not encode a priori assumptions that could inform a lab manager or be used for inference. The probabilistic error semantics from Oops could be integrated into these systems’ protocol interfaces, which are often no-code.

Although Aquarium and Benchling are intended more for organization than prediction, Antha is capable of executing protocols *in silico*, meaning through digital simulation. With a model of underlying processes, integrating Oops into Antha could enable both error inference and fitting error parameters such as probabilities and uncertainties to real execution traces.

Puddle [26] can autonomously recover from errors made when executing protocols on digital microfluidic devices. By detecting when droplet positioning diverges from what is expected, Puddle synthesizes instructions to reset to an error-free state. Microfluidic automation systems like Puddle could execute subprotocols in Oops and eliminate sources of error for Oops to worry about.

### Metaprogramming and Domain-Specific Languages

Oops builds on previous metaprogramming languages and domain-specific languages to achieve concise and flexible specification of biological protocols. Much like prior generative metaprogramming systems [11, 12, 16, 19], Oops programs include metaprograms that transform ASTs to reduce clutter in the original programs and automate repeated rewrites. The Oops language is designed around the lab as an unconventional computing platform and allows exploration of protocol design. Recent work has explored languages for exploring the design space of other unconventional computing methods such as analog circuits in Ark [25].

## 8 Limitations & Future Work

### Executable Protocols

Oops protocols are not executable, because they do not encode all the information necessary to be physically run by humans or robots. One action in Oops can map to multiple real lab operations. One protocol represents what can happen to a single sample, whereas a real protocol batches together many samples to execute operations on simultaneously. Oops could be lowered to a finer-grained protocol language used by an existing biofoundry. Implementing a system like Oops in a biofoundry would enable a feedback loop between the model and the real-world process.

### Cost Optimization

Failing fast (Section 6.4) is only one metric to explore the design space of protocols. With a formalized model of how a protocol can fail, attaching costs (in time or money) to each action would unlock simulations of the cost to achieve successful outcomes and variance in product. By searching over the space of protocols with different controls and verification steps enabled, Oops could optimize costs.

## Footnotes

1 The terse variable naming is intended to reflect lab naming conventions.

